# The role of learning in the evolution of status signalling: a modeling approach

**DOI:** 10.1101/2023.09.19.558427

**Authors:** Andrés E. Quiñones, Redouan Bshary, C. Daniel Cadena

**Author notes:** Correspondence: Andrés E. Quiñones < >.

## Abstract

Adaptive behavioural responses often depend on qualities of the interacting partner of an individual. For example, when competing for resources, an individual might be better off escalating fights with individuals of lower quality, while restraining from fighting individuals of higher quality. Communication systems involving signals of quality allow individuals to reduce uncertainty regarding the fighting ability of their partners and make more adaptive behavioral decisions. However, dishonest individuals can destabilize such communications systems. An open question is whether cognitive mechanisms, such as learning, can maintain the honesty of signals, thus favoring their evolutionary stability. We present evolutionary simulations where individuals can produce a signal proportional to their quality and learn along their lifetime the best response to the signals emitted by their peers. Our simulations replicate previous results where the handicap principle mediates the evolution of signals. In our simulations learning on the receiver side can mediate the evolution of signals of quality on the sender side. When the cost of the signal is proportional to the quality of the sender, all individuals in populations are honest signalers. In contrast to traditional models which predict the absence of signals when the cost is not proportional to the quality of the signaler, our model revealed that learning facilitates the evolution of a polymorphism in which populations comprise both honest and dishonest signalers. We argue that learning may have a role in the evolution and dynamics of a wide range of communication systems and more generally in behavioral responses.

## Introduction

The outcome of interactions with conspecific individuals is a critical determinant of fitness in social animals. Irrespective of whether such interactions are cooperative or competitive, the actions of interacting individuals influence their reproductive success. However, the best action for an individual engaging in a social interaction might vary depending on its own condition and that of individuals with whom it interacts. Thus, acting based on information about interacting partners is typically adaptive (Quiñones et al. 2016), yet acquiring such information is far from trivial. In some cases, interacting partners (*i.e.* signallers) might be ‘willing’ to provide accurate information, but in others it might be in their own interest to conceal information (Johnstone 1997) or to provide deceptive information (Johnstone and Norris 1993). Typically, in a given context, some individuals benefit may from broadcasting accurate information, whereas others may benefit from concealing it. Take, for instance, an interaction between two individuals where one can help the other. Given some costs and benefits, a potential donor is likely to be interested in helping related individuals. Therefore, for a relative of the donor, broadcasting kinship would be advantageous, whereas concealing the lack of kinship would be better for an unrelated individual. Similar scenarios may apply to a variety of aspects of social life such as finding mates, feeding offspring, or engaging in dominance relationships or aggressive interactions (Bradbury and Vehrencamp 2011; Møller 1988; Tibbetts and Dale 2004).

In dominance and aggressive interactions, a crucial piece of information to guide individual actions is the fighting ability of interacting partners, which is sometimes referred to as Resource Holding Potential (RHP, Parker 1974) or simply as quality. Responsiveness to the quality of an opponent is central to communication systems in which individuals exhibit badges of status, where a signal typically conveys quality (Johnstone and Norris 1993; Rohwer 1975). Such signals tend to be arbitrary, however, meaning the signals could potentially be produced by high- and low-quality individuals alike, opening up the possibility of dishonest signalling and raising the question of evolutionary stability. Signals can be evolutionarily stable whenever low-quality individuals pay a higher fitness cost for carrying the signal such that it is not in the interest of low-quality individuals to fake quality by producing a large badge. Signal costs inversely proportional to quality may come about as a result of the production costs of the signal, in which case the signal works like a handicap (Botero et al. 2010; Johnstone and Norris 1993; Grafen 1990; Zahavi 1975). Alternatively, the costs can come from the aggressive reaction of receivers when the convention of the communication is broken (Enquist, Ghirlanda, and Hurd 2010; Tibbetts and Dale 2004). For example in paper wasps, subordinate individuals with experimentally manipulated dominant-like facial patterns received more aggression from the dominant (Tibbetts and Dale 2004). In this case, costs are socially rather than developmentally mediated and are triggered by the violation of the convention, although how the convention is established in the first place is unclear.

A somewhat ignored component of communication systems in the context of aggressive interactions are cognitive aspects of the receiver module. Theoretical models often assume that communication systems relying on badges of status have reaction norms as their mechanistic underpinning (Botero et al. 2010), with individuals using their own quality and the badges of opponents to determine whether to aggressively engage in contests. Reaction norms allow individuals to respond to the available information without a large cognitive burden because the computations involved in the reaction norm response do not require much memory nor advanced algorithmic processes. This contrasts with other information-processing mechanisms like individual recognition where individuals associate cues of their peers to their quality. Because these associations must be learnt throughout the life of individuals, they have the usual cognitive requirements of associative learning processes. An alternative view of badges of status is that individuals learn to react to them based on their experiences (Guilford and Dawkins 1991), which would imply that receivers must learn to associate signals with the fighting ability of bearers just as in systems based on individual recognition. In cases where badges of status vary quantitatively (*e.g.* in size or intensity), fighting ability may increase monotonically with attributes of the badge and would be reinforced by every interaction. Therefore, in principle, learning in such scenarios would be faster than in systems involving individual recognition in which the association between signals and their meaning varies depending on the interacting partners. In any case, learning may be a central cognitive mechanism in both types of communication systems, but the role of learning in these contexts has not been thoroughly explored in the empirical nor theoretical literature.

Associative learning is a taxonomically widespread cognitive mechanism that allows individuals to associate rewards with environmental stimuli and thus behave adaptively (Heyes 2012; Macphail 1982; Staddon 2016; Behrens et al. 2008). Theory has shown that natural selection favours such associations in complex environments where conditions are difficult to predict (Dridi and Lehmann 2016). Associative learning is a flexible cognitive mechanism whose underpinnings show interspecific variation (Enquist, Lind, and Ghirlanda 2016; Quiñones et al. 2020; Prétôt, Bshary, and Brosnan 2016; Prétôt et al. 2021). For example, species can have different learning rates (Hoedjes et al. 2010), some may vary these rates during learning trials which allows them to have more flexible learning (Leimar, Quiñones, and Bshary 2023), and some may include future rewards in their associations (Raby et al. 2007; Osvath 2009). Associative learning is not a single mechanism, but rather a set of cognitive structures and processes that can vary in their scope and complexity. Presumably, these structures and processes have been modified by natural selection and could provide novel explanations for behavioural variation. Nonetheless, associative learning is not often included in the narrative of evolutionary explanations of behavioural patterns (Fawcett, Hamblin, and Giraldeau 2013; Kamil 1983; McAuliffe and Thornton 2015).

Computational models of evolution can overcome the lack of integration between learning and evolution. Reinforcement learning theory encompasses a series of computational methods inspired on the psychological and neurological mechanisms of associative learning (Sutton and Barto 2018). This set of algorithms allows the implementation of biologically realistic problems, capturing the essence of learning processes (Frankenhuis, Panchanathan, and Barto 2018; Quiñones et al. 2020). Furthermore, these algorithms can be embedded in evolutionary simulations to generate theoretical predictions of the effect of learning on behavioural evolution (Leimar and McNamara 2019; Leimar and Bshary 2022b).

Here, we present an evolutionary model where individuals use associative learning to develop a tendency to behave either aggressively or peacefully in the context of competition over resources, depending on a quantitative morphological trait (*i.e.* a badge indicating quality) they perceive in their opponents. Over evolutionary time, the size of the badge evolves as does its dependency on the individual’s quality. Under this simple set up, individuals can use the badge as a signal of quality. We use the model to assess under what conditions one might expect communication signals to evolve as handicaps or conventions.

## The model

We model the evolution of signals indicating individual quality in the context of agonistic interactions. For simplicity, we consider a population of haploid individuals with non-overlapping generations. Individuals are born every generation with a level of quality (*Q*_*i*_, where *i* is subscript of the population vector of size *N*) given by a number between zero and one, which is drawn from a truncated normal distribution *N*(0.5, σ). An individual with quality 0 has the lowest RHP, while an individual with quality 1 has the highest. As they develop, individuals produce a phenotypic signal (badge), the size of which depends on their quality according to a reaction norm given by ***B***_*i*_ = 1/(1 + *e*^α*i*−β*iQi*^); where α_*i*_and β_*i*_are individual specific traits that determine the shape of the reaction norm (Fig. 1 A). Variation in α_*i*_ and β_*i*_ means individuals can have either uninformative (flat) or informative (logistic) norms. Informative reaction norms represent a developmental program where the phenotype of the individual is determined by the environmental conditions under which it grows, thus causing a correlation between quality and signal. The size of the signal is constrained to take values between zero and one. We assume different values of these traits are given by different alleles and are inherited from mother to offspring unless, with a small probability (μ), mutation changes the allelic value of the offspring by an amount drawn from a normal distribution *T N*(0, σ_μ_).

**Figure 1:**
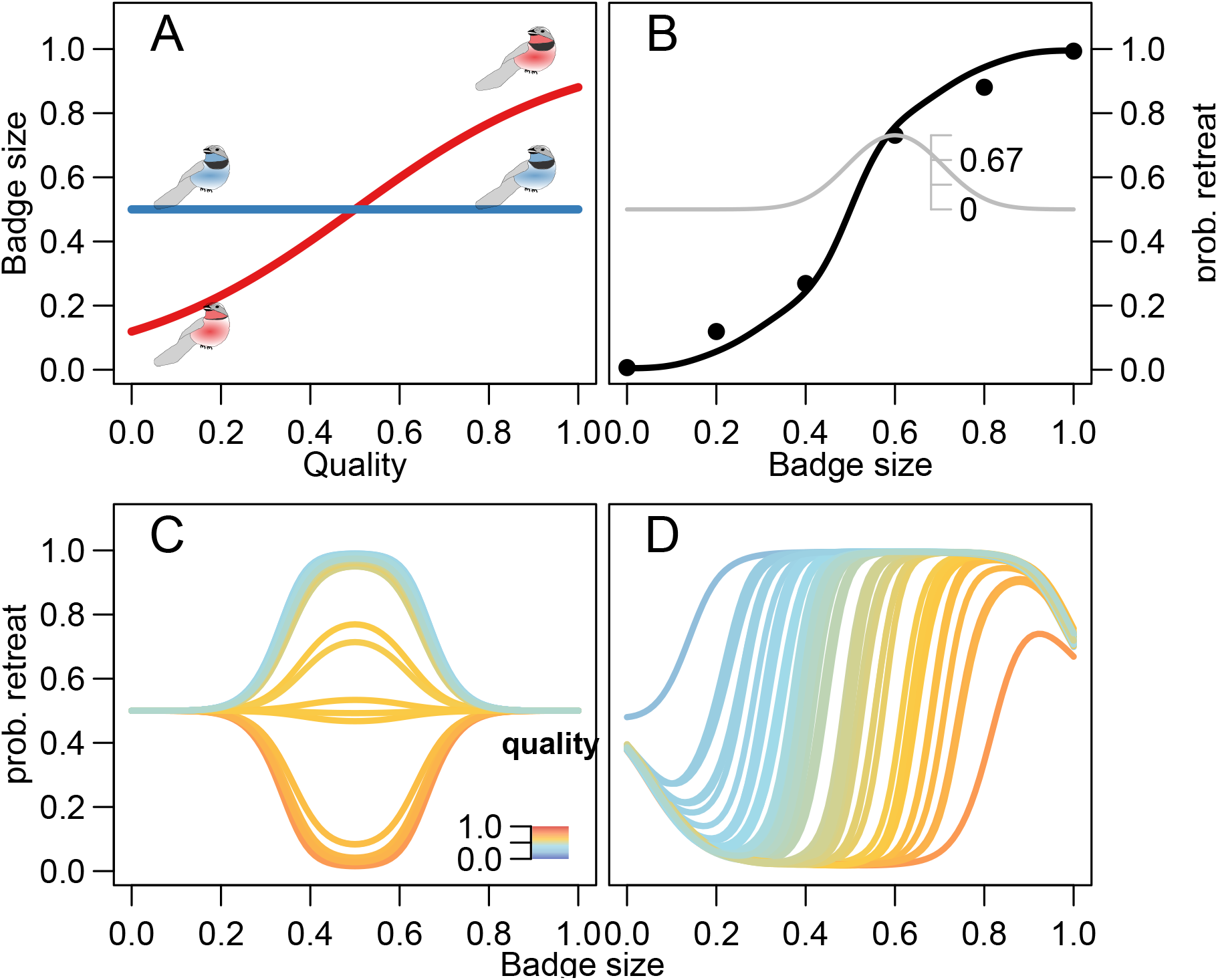
Model of communication in the context of aggressive interactions. In A, the reaction norm determines the badge size, *e.g* the size of a black band on a bird’s chest. Red shows an informative reaction norm, whereas blue shows a uninformative reaction norm. B individuals perceiving the signal have a behavioral reaction norm that determines their probability of retreating (black line). The reaction norm arises from generalizing the information from the feature weights (black dots). Feature weights are values associated with a particular badge size and are updated as individuals interact with each other. The generalization of the information associated with each feature weight is represented by the grey line and its axis, which shows that the response triggered by the fourth feature diminishes as the value evaluated is further from the feature center. The learning process moves the feature weights (black dots) up or down depending on whether this leads to an increase in the estimated reward. Panels C and D show the effect of learning on the receiver strategy. Receivers in C face signallers with uninformative reaction norms (blue line in A). The uninformative reaction norm yields a badge size of 0.5, thus the behavioural reaction norm in C is only updated around that value. D Receivers face signallers with informative reaction norms (red line in A). Accordingly, receivers develop a threashold-like reaction norm where the decision to retreat increases with the badge size of the interacting partner. Colour scale in C and D indicates the quality of the individual.

After birth, individuals go through a round of viability selection. The survival probability of an individual is given by *s*_*i*_ = 1/(1 + *e*^−*k*1−*k*2(*Qi*−***B****i*)^), where *k*_1_and *k*_2_ are parameter values determining the shape of the survival function. Importantly, if *k*_2_ = 0 survival probability is independent of quality, while if *k*_2_ > 0 lower-quality individuals pay a higher price for similar-sized badges, fulfilling the core assumption of the handicap principle (Botero et al. 2010; Grafen 1990; Johnstone and Norris 1993).

Individuals who survive engage in a series of pairwise interactions where they compete for resources. In each interaction individuals must decide whether to escalate a fight or not. Following the classic *hawk-dove* game (Maynard-Smith 1982), if the focal individual fights and its partner does not, then the focal gets as pay-off the resource of value *V* while its partner gets nothing; if both individuals restrain from fighting, then they split up the resource in half. If both individuals decide to fight, then the winner takes over the resource and the cost of fighting is split between the two players. We further assume that the probability of wining a fight for the focal individual depends on the difference in quality between it and its interacting partner; specifically it is given by *p*_*ij*_ = 1/(1 + *e*^−*k*3(*Qi*−*Qj*)^), where *k*_3_ is a parameter defining how strong the quality difference determines the winning probability, and *i* and *j* denote the the position in the population vector of the focal and its interacting partner, respectively.

The decision of whether to escalate a fight against an interacting partner can depend on the size of the partner’s badge, with the dependency being determined by the focal’s experiences acquired through a learning process. We implement learning using the actor-critic approach from reinforcement learning (RL) theory (Sutton and Barto 2018; Quiñones et al. 2020; Leimar and McNamara 2019; Leimar 2021). Individuals estimate the reward (pay-off) expected from interacting with partners of different badge sizes (the critic in RL terminology). After each interaction they update the estimate of reward proportionally to the difference between their current estimate and the observed reward (prediction error δ) and to the speed of learning (A). Furthermore, individuals express different probabilities of retreating/attacking depending on the badge size of their opponent (the actor in RL). They update the probability of retreating/attacking depending on whether retreating/attacking leads to an increase in the reward estimation. Thus, if a focal individual decides to escalate a fight against an individual with a small badge and this leads to an increase in the reward estimation, then the focal individual will increase the probability of escalating fights with individuals of small badges in the future. Given that badge size is a real number between 0 and 1, there are infinitely many badge sizes. Thus, the reward estimation, as well as the probability of retreating/attacking, must be generalized across different values. To implement generalization we use the linear function approximation method based on radial basis functions (Sutton and Barto 2018). Specifically, we pick *c* feature centres, which are evenly spaced values along the badge size interval ([0,1]) where the updates are focused. For simplicity, we keep the location of these feature centres constant and the same for all individuals; they are stored in vector **b**. Each of these feature centres is associated with a weight for reward estimation and tendency to play retreat in a given interaction. The reward estimation and the tendency to retreat are calculated (in every interaction) as the sum of the weights associated with each feature centre, weighted by the response triggered by the feature (Fig. 1 B, dots represent the feature weights). The response of each feature centre diminishes as a Gaussian function with the distance between the feature centre and the badge size of the partner (Fig. 1 B, grey line).

Formally, the reward estimation 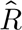when the focal individual (*i*) faces individual *j* is given by,

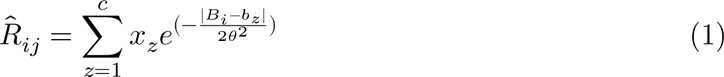

where *x*_*z*_ is the weight of feature centre *z* on the reward estimation; and θ is the width of the generalization function. Similarly, the tendency to retreat is given by the sum of feature weights associated with the actor, and the probability is obtained by applying a logistic transformation (Fig. 1 B, black line). Thus formally, the log-odds to retreat when facing individual *j* is

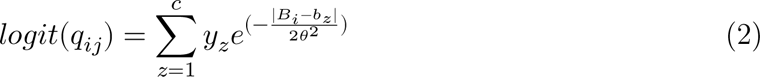

where *y*_*z*_ is the weight of feature centre *z* on the tendency to retreat.

Individuals interact as the focal individual *n* times and the interaction partner is chosen at random. Thus, the expected number of interactions for each individual is 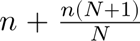. After the interaction round, individuals in the population reproduce with a probability proportional to their total pay-off *w*_*i*_, which is a sum of the baseline pay-off (*w*_0_) and all the pay-offs obtained throughout their life. Thus, the combination of natural selection and genetic drift changes the distribution of values in α and β that segregates in the population, effectively changing the badges expressed and the communication system.

The code necessary to run simulations as well as to produce visualizations can be found in: https://doi.org/10.5281/zenodo.8359809

## Results

### Learning in a monomorphic population

We first present the outcome of simulations where we prevented the evolution of the badge size (by setting the mutation rate to 0), and assumed that all individuals in the population display either an uninformative or informative badge size with respect to their quality. These simulations show what type of receiver strategy develops through a learning process in such a monomorphic population. Figure 1 (C and D) shows the receiver strategy developed through learning; panel C is for receivers that faced uninformative signals, and D for those facing informative ones. The simulations reveal that when learners face uninformative signals (Fig. 1 C), they modify their probability of retreating depending on their own quality. Individuals with higher quality (red tones) have a high probability of attacking after the learning process while individuals of lower quality (blue tones) mostly retreat from confrontations; in both cases, individuals behave equally irrespective of the quality of their opponent. Thus, learning splits the population of receivers into the two classic pure strategies of hawks and doves. Given that we assumed a monomorphic population with unresponsive reaction norms on the signalling side, the changes triggered by learning only affected a small range of badge sizes. Specifically, all learning occurred around a badge size of 0.5, because all badges in the population are of this size (panel C). In contrast, when receivers face informative reaction norms on the side of the signaller (Fig. 1 B), receivers use information on the badge size of their interacting partners to determine whether to retreat or attack. The relationship is given by a threshold-like reaction norm, where the decision to retreat or to attack depends on the quality of the receiver. As expected, the higher the quality of the receiver, the larger the badge size of the signaller that triggers a retreat.

### Badges as Handicaps

When we allowed the badge size to evolve (α and β changed subject to natural selection and genetic drift) and the signal worked as a handicap (*i.e.* the cost of the badge was inversely proportional to the quality of the individual), the sender code often evolved to produce an honest signal (Fig. 2). The evolution of the sender code did not happen immediately after the start of the evolutionary process. Instead, the evolutionary dynamics of the reaction norm parameters (Fig. 2, A) appeared to involve a set of steps. First, the intercept of the reaction norm (α) describing the relationship between individual quality and badge size evolved to higher values, reducing the average badge size in the populations. This is because reaction norms segregating in the population are flat at the beginning of the simulation; consequently, the receiver code does not respond to the badge size and larger badges are costlier and do not trigger lower attack probabilities. During the first generations, the slope of the sender reaction norm (β) remains close to the initial value of zero. At around generation 4000, the slope evolves toward positive values. Positive values in the slope imply that larger badges correlate with higher quality (Fig. 2 B). Receivers then learn to react to such correlation, increasing the probability of retreating when facing individuals with larger badges (Fig. 2 C). Hence, natural selection favours larger values of β, eventually leading to an evolutionary equilibrium in which badge size is an honest signal of quality mediated by the learned responses of receivers (Fig. 2 B.4 and C.4).

**Figure 2:**
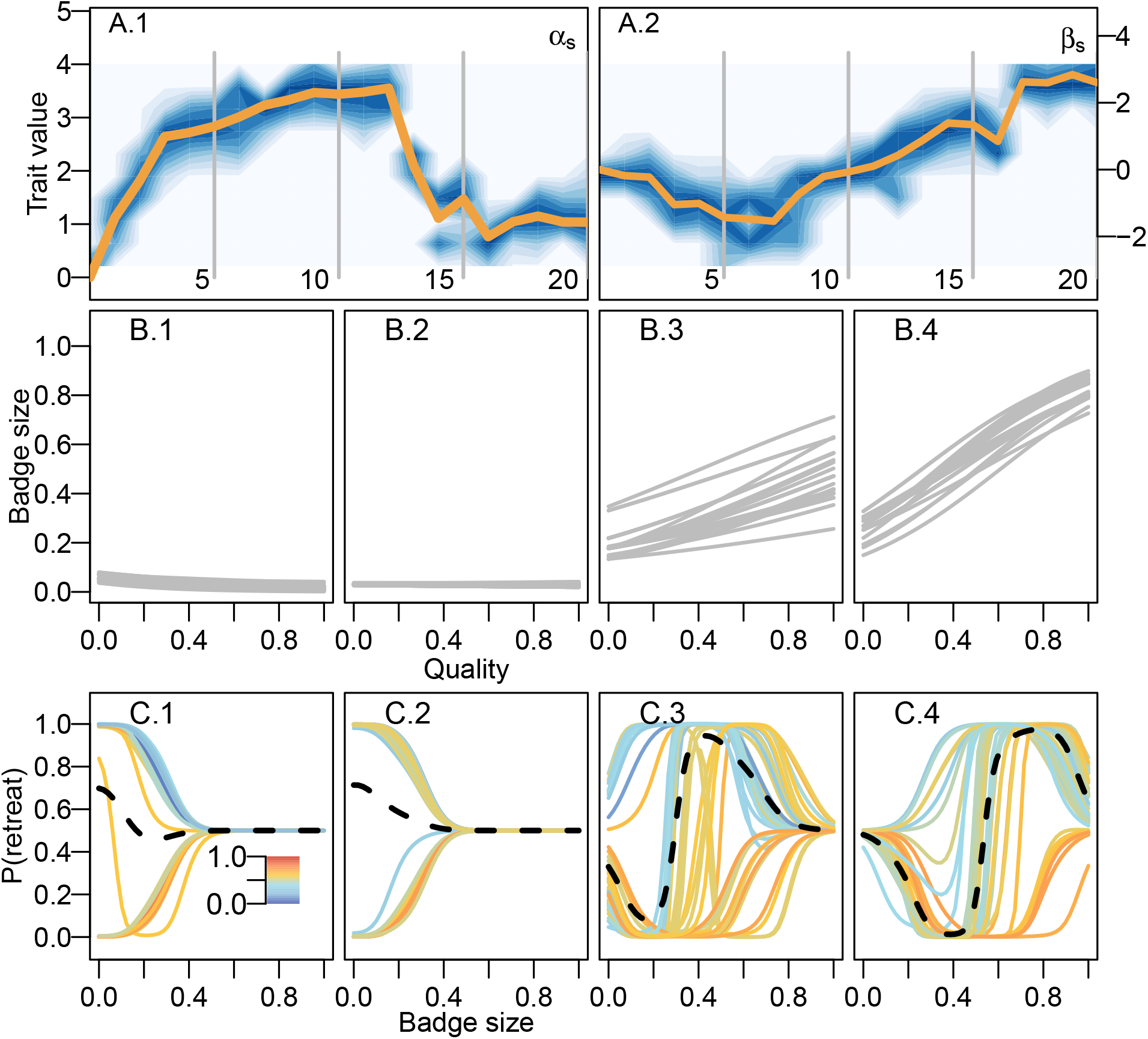
Evolution of the badge size as a handicap mediated by learning. Top panels (A.1 and A.2) show the evolutionary dynamics of the sender code. On the left (A.1), changes in the distribution of values for the intercept of the reaction norm (α); on the right (A.2) changes in the distribution of the slope (β). Darker areas of the background correspond to values with high frequency, while orange lines in both panels show the mean of the distribution. X axis in the evolutionary dynamics are given in thousands of generations. Grey lines show the generation time corresponding to the panels below portraing the sender (B) and receiver code(C). In the bottom panels (C1-4), the learned reaction norms correspond to individuals in the population after the interaction round. Color scale indicates quality just as in Fig. 1. On the first half of the simulation (B.1 and B.2) the badge size evolves to its minimum value. In the second half, the slope takes positive values and reaction norms evolve to produce an honest signal of quality. Correspondingly, individuals learn to react to badge size by increasing the probability of retreating with increasing badge sizes.

The evolutionary trajectory portrayed in figure 2 A is not the only possible outcome. If the slope of the sender reaction norms (β) evolves, subject to genetic drift, toward negative values before badges become handicaps, receivers never learn to react to the size of badges. Thus, badges are only costly and do not provide information about the quality of individuals (Fig. S1). Eventually, therefore, badges disappear from the population. In contrast, when the badge does not work as a handicap but instead is cost-free, the evolutionary process never leads to the establishment of an honest signal. Instead, subject to genetic drift, the population either evolves towards the disappearance of the badge or to the maximum badge size. In either of these cases reaction norms are flat, so the badge does not provide any information about quality.

### The evolution of polymorphism mediated by learning

The amount of information that agents are able to collect via learning over their life time can strongly change the outcome of evolutionary dynamics. In the simulations presented in figure 2 and S1, individuals learned with high speed (A = 0.4) and interacted repeatedly along their lifetime (2000 interactions on average). When we reduced the number of interactions that individuals experienced over their lifetime to 300 on average, we saw a drastic increase in the phenotypic and genetic variation present in the population. Populations start monomorphic with a value of zero for both the intercept and the slope of the reaction norm, and mutations quickly build up a normal distribution around the starting value. Within the first 2000 generations, the unimodal distribution in the intercept (α) splits into a bimodal one. Later in evolutionary time, one of the peaks splits further, so at the end of the evolutionary simulation the distribution of the intercept (α) in the population shows three distinct peaks. In the case of β, the peaks in the distribution are not so clear-cut, but the variance of the distribution increases over time. These changes in the distribution of the parameters of the sender reaction norm imply that individuals can generally be classified into three distinct types (Fig. 3). Two types express a flat reaction norm with extreme values for the badge size, meaning that their badge size is not informative of their quality, whereas the third type shows intermediate badge sizes determined by individual quality (Fig. 3 B.3-4). In this simulation, the values of the slope in the informative group are negative, which implies that badge size correlates negatively with quality. Furthermore, the receiver reaction norms developed through learning, particularly those of individuals of intermediate quality, respond to the signal of their sender type by increasing the probability of retreating from a fight with individuals with smaller badges (Fig. 3).

**Figure 3:**
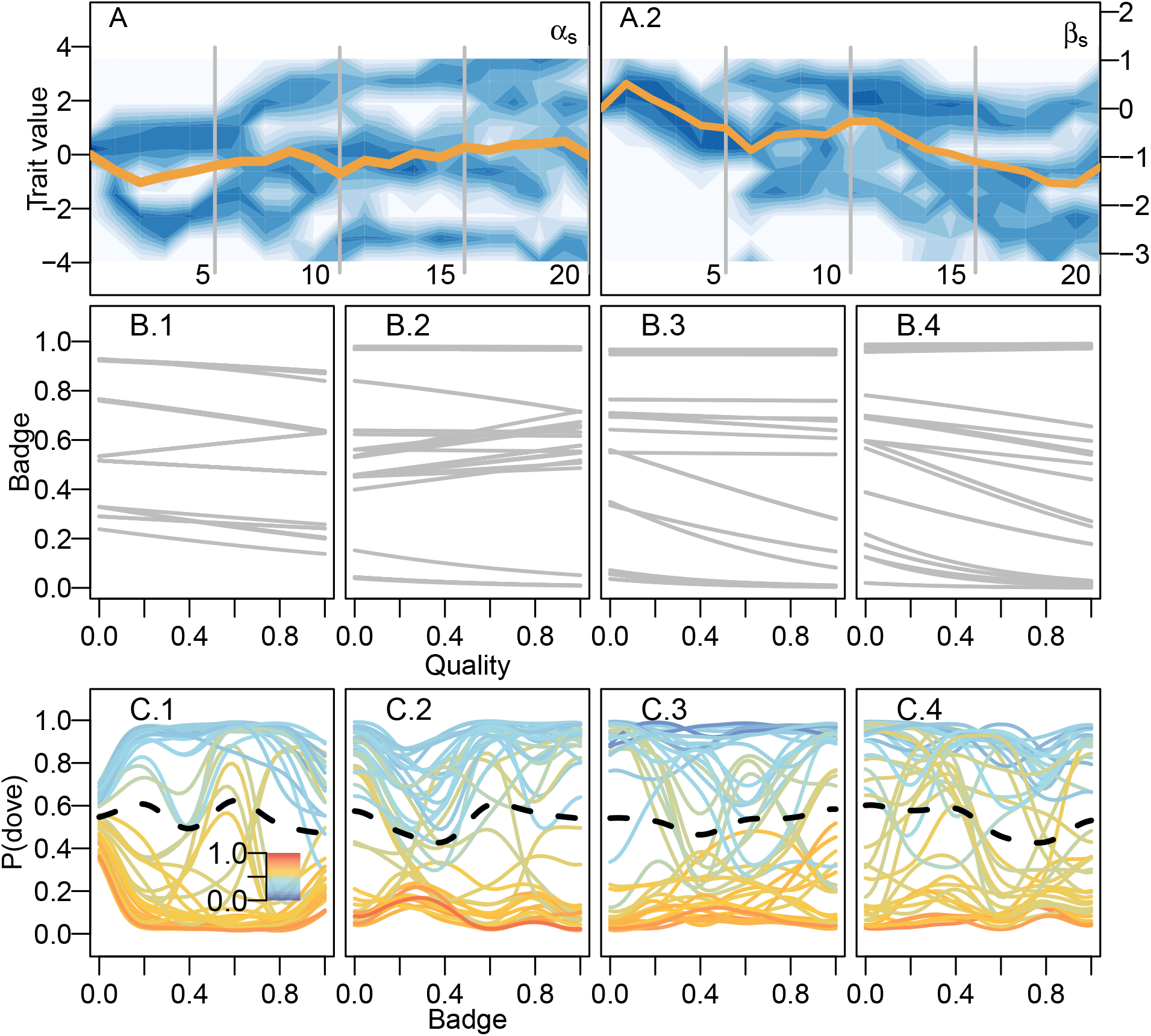
The evolution of cheap signals. Portrait of the evolutionary dynamics of the sender code with snapsots of both sender and receiver codes just as in fig. 2. The top panels (A) show changes in the distribution of values for α (A.1) and β (A.2) along evolutionary time. Panels below correspond to snapshots of the sender (B) and receiver codes (C). Generation time of the snapshots are indicated by the grey lines in the top panels. The unimodal distribution with which α (A.1) starts the simulation, quickly turns into a bimondal distribution, and at about 10000 generations it splits further into three modes. These three peaks correspond to three type of reaction norms in panels B.1-3.

A limited number of interactions has an effect on the amount of variation in the evolving parameters when the signal follows the handicap principle as well. In fig. 4, we show simulations where individuals have on average 300 interactions in their life and the cost of the signal is proportional to quality, following the handicap principle. The evolutionary process leads to a combination of reaction norm parameters where there is a positive correlation between the size of the badge and the quality of the individual. This relationship, however, is muddled by the fact that there are two clusters of values for the intercept (α) and slope (β) of the reaction norm in the population. Thus, there are two types of reaction norms. One of the types has a steeper slope compared to the other. This effect of an increased variance in the trait distribution is not only triggered by lower number of interactions. Larger variances are also found when we assume a lower speed of learning (data not shown). This suggest that limits to the amount of information that individuals acquire through learning allows the coexistence of different communication strategies within a population, something previously shown by Botero et al. (2010).

**Figure 4:**
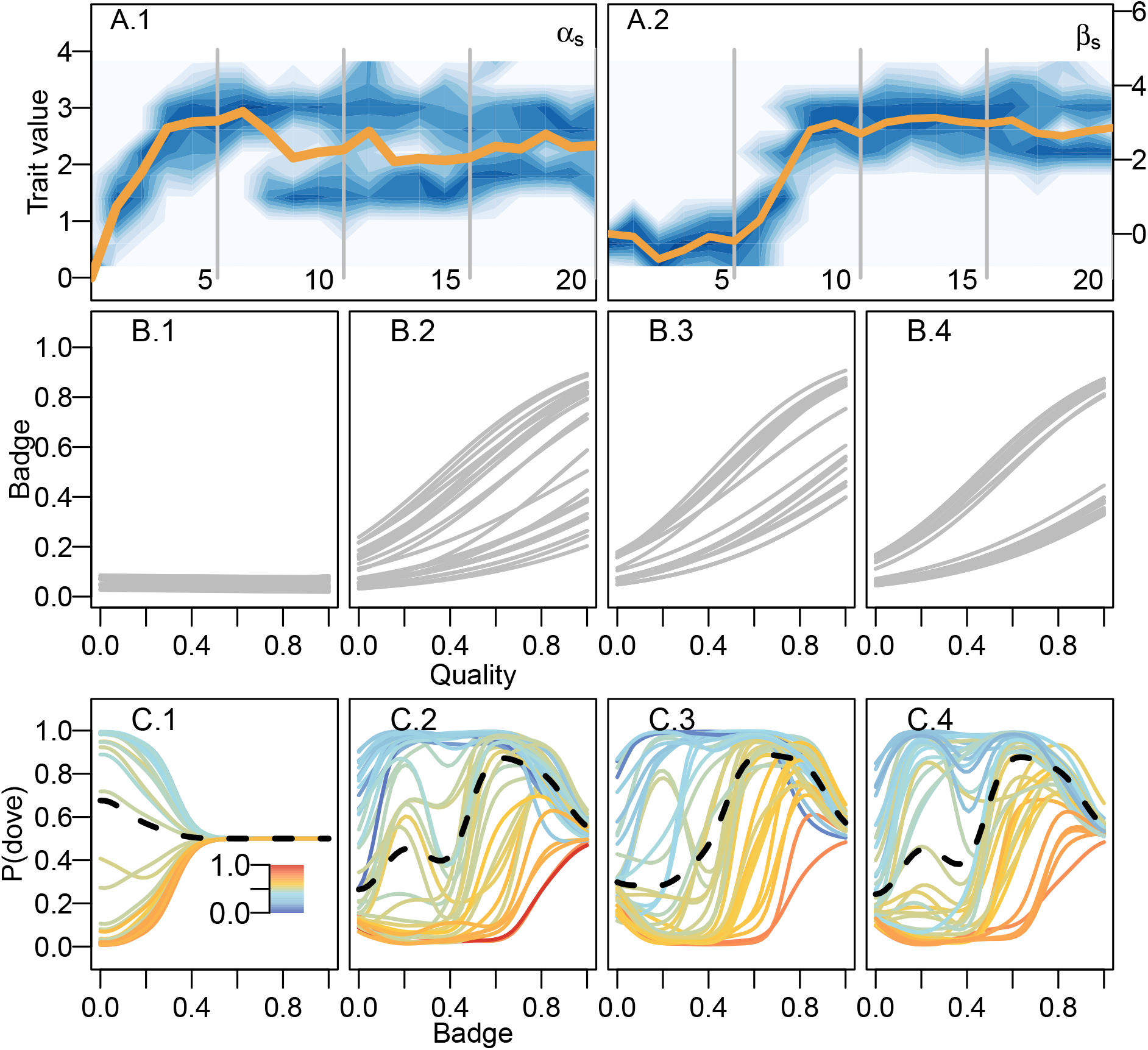
The evolution of costly signals. Portrait of the evolutionary dynamics of the sender code with snapsots of both sender and receiver codes just as in fig. 2. The middle panels show changes in the distribution of values for α and β along evolutionary time. Panels above and below correspond to snapshots of the sender and receiver codes, respectively, generation time of the snapshots are indicated by the grey lines in the middle panels. Similarly to the evolutionary dynamics in Fig. 2, first the badge size evolves towards its minimum value. Then, the slope evolves positive values determining an increasing reaction norm. In contrast to 2, the normal distribution splits up into two modes. This translates into two types of reaction norms assorting in the population where both code for a positive relation between quality and badge size.

### The peaceful, the aggressive and the clever

Our model revealed that the behaviour expressed by naive individuals (*i.e.* those who have not yet learned) imposes negative-frequency dependent selection, allowing for the build up of genetic variation in reaction norms. In the simulations presented so far, we assumed that individuals start with a flat behavioural reaction norm such that they escalate fights aggressively with a 0.5 probability regardless of the badge size of the interacting partner. To assess whether such initial conditions of the communication system had any role in the build up of genetic variation, we ran a series of simulations varying the initial conditions of the actor module in the learning model. Specifically, we let naive individuals have a flat reaction norm with either a 1) low (peaceful), 2) high (aggressive) probability of escalating a fight or 3) a probability corresponding to the ESS of the original hawk-dove game (which we call clever, due to information and computation necessary to know the ESS). Fig. 5 shows the distribution of values of the intercept (α) and slope (β) evolved in different replicates of the simulations. The left-hand side panel, corresponding to peaceful naive individuals, is the only one where the distribution of values is split in different clusters. That is, most of the variation occurs within clusters. In contrast, when naive individuals behave either aggressively or cleverly, badges evolve toward minimum and maximum values, and so the variation is driven mainly by differences among replicates. We can make sense of these results by realizing that individuals change their naive behaviour quickly in ranges of the badge size that are common in the population. In contrast, an individual expressing a rare badge size will most likely experience the naive behaviour of their partner. When the naive behaviour is peaceful, individuals with a rare badge size have a fitness advantage. This triggers negative-frequency dependent selection and the evolution of different types of badge sizes. According to this narrative, situations where individuals learn fast and interact repeatedly will diminish the strength of frequency-dependent selection.

**Figure 5:**
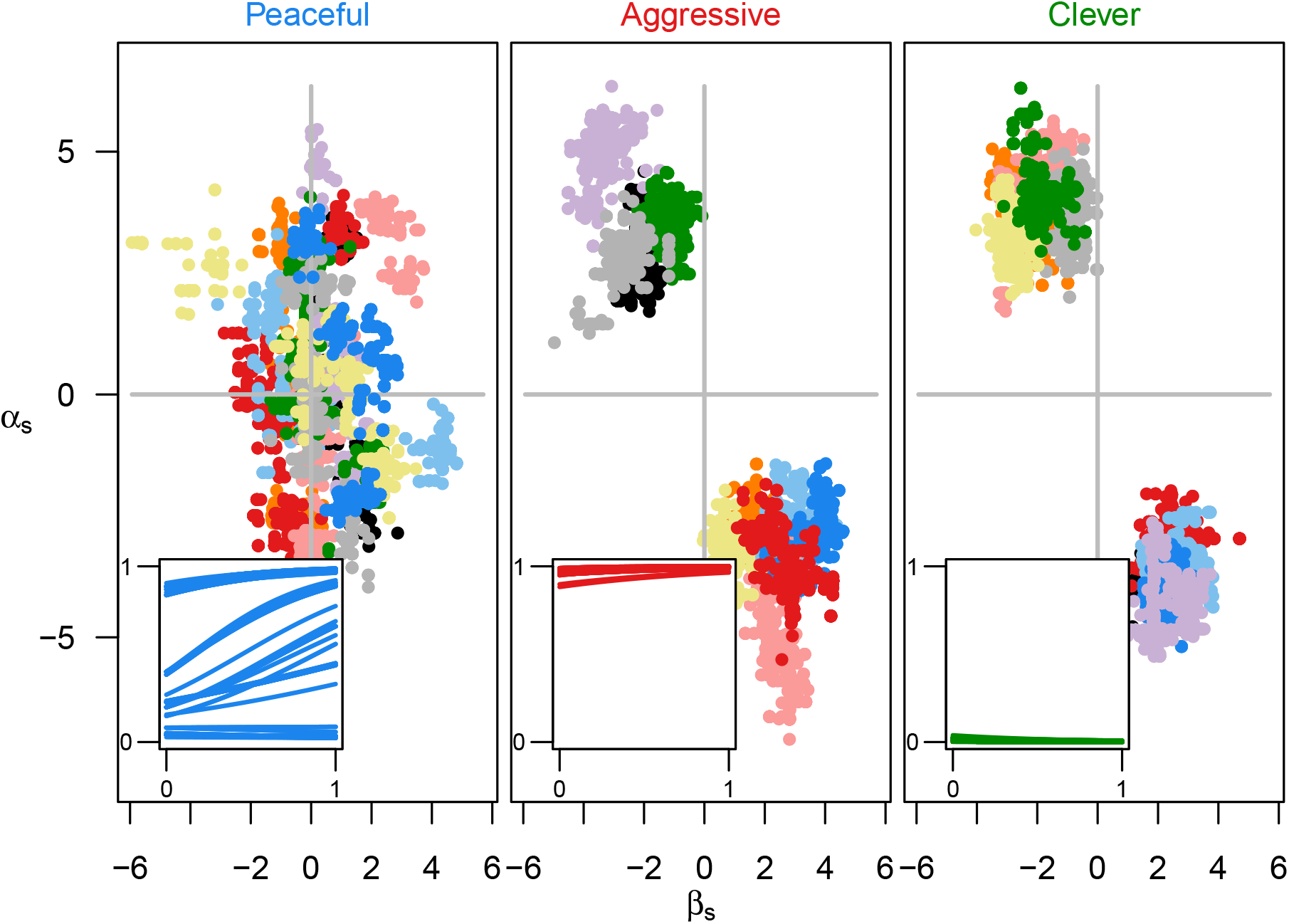
The peaceful, the aggressive and the clever. Distribution of values of the intercept α and slope β for individuals at the end of the evolutionary simulations. The panels show the three different initial conditions for the behaviour of individuals: peaceful, agressive and “clever”, see maintext for details. Colours indicate the replicate simulation. In the inset, the resulting reaction norms corresponding to the intecept and slope values for one of the replicates, which replicate is shown is indicated by the line colour. The simulations where individual disply a “clever” and agressive behaviour, before the learning process, show all the points of a single replicate cluster together, while in simulation where individuals start being peaceful al replicates show distinct groups of individuals spread throughout the x and y axis. Only in this condition there is a frequency-dependent process that favour diversity in the reaction norm.

The action of natural selection and genetic drift on the learning parameters do not override the conditions for the evolutionary branching (Dieckmann and Law 1996) described above. Given how crucial naive behaviour is for the branching event described earlier, we tested whether evolutionary processes could lead key parameters of the learning process toward values that prevented the genetic variation to build up. Figure S2 shows the dynamics from a set of simulations where we allow for the coevolution of the reaction norm as well as key parameters of the learning module, namely the speed of learning (*A*) and the behavioural tendency expressed by individuals prior to learning. Top panels in figure S2 show that there is diversification process in this coevolutionary scenario as well. That is because, as is shown in the middle panels (B) of the same figure, the learning parameters stay within the range necessary for the branching event to take place. The speed of learning (Fig. S2 B.1) stays well above zero throughout the simulation. In contrast, the initial behavioural tendency maintain values around zero. Thus, evolutionary processes acting on the learning parameters did not override the conditions necessary for frequency-dependent selection to boost genetic and phenotypic variation.

## Discussion

We presented a model that integrates learning with evolutionary processes. In the model, the response to a signal in a communication system is mediated by learning processes: individuals learn how to respond to a quantitative trait of their interactive partners. We showed that the learning process can mediate the evolution of an honest signal under the handicap principle (Grafen 1990; Zahavi 1975). This conclusion is not different from classical genetic models, where responses to signals are innate to the individual. However, unlike classical genetic models, our model shows that learning can also mediate the origin of a signal polymorphism in the absence of the handicap principle. Under this polymorphic equilibrium the population consist of distinct types of individuals, with honest and dishonest signals. The amount of information gathered through learning, as a well as the initial conditions of the learning process, were crucial for the emergence and maintenance of variation in the signalling system. Namely the polymorphism required a limit in the amount of information collected by individuals, as well as peaceful behaviour in naive individuals. Furthermore, we showed that the conditions leading to the origin of the polymorphism can be reached when evolutionary processes are able to change the parameters of the learning system (the speed of learning and the naive behaviour).

Associative learning is a powerful mechanism to learn about the world, about social partners, and about one’s abilities in a particular social context. By associating cues and signals with fitness-relevant outcomes, individuals collect information allowing them to improve their reproductive potential. This is evident in classical situations such as when animals learn to avoid food that makes them sick or use environmental cues to find food. More recent theoretical work has highlighted the potential role that associations can have in social contexts such as hierarchy formation (Leimar 2021; Leimar and Bshary 2022a, 2022b) and cooperation (Leimar and McNamara 2019; Dridi and Akçay 2018). In these examples, models involve individuals who use various sources of reward to make adaptive choices. Here we extended this logic to the evolution of a communication system mediated by a signal of quality. Previous evolutionary models of badges of status assumed that individuals responded using a behavioural reaction norm, where the opponent’s badge and the individual’s own quality determined behaviour (Botero et al. 2010). However, there is no clear mechanism justifying the assumption that individuals inherently know their own quality, particularly relative to their peers. In our model, individuals not only learn about the quality signal, but also learn about their own quality relative to others. This can be seen in our simulations in the responses developed by individuals of different quality. Because individuals learn about their own quality, even in the absence of an honest signal, they are able to make more adaptive choices. Specifically, lower quality individuals refrained more from escalating fights, while higher-quality individuals took over resources they could win in a fight. Thus, the learning process we modelled and the response it mediates has fitness-relevant consequences even in the absence of an honest signal. This was confirmed by simulations where the speed of learning was allowed to change subject to evolutionary processes. In those simulations, natural selection always maintained learning rates above zero. Our simulations however, only captured the effect of selection mediated by the social game on the learning parameters. For example, we did not account for potential costs of faster updating, or improved performance in foraging tasks. Inter-specific variation in cognitive abilities mediated by environmental differences has been reported elsewhere (Sonnenberg et al. 2019). If such variation is partly mediated by changes that affect general cognitive processes, like the speed of learning, they could impact the outcome of communication systems like the one we modelled.

Learning processes which involve collecting information on a population-wide level promote frequency-dependent selection. For example, frequency-dependence triggered by learning process was highlighted by a classic study where live predators drive the evolution of *in silico* polymorphic prey (Bond and Kamil 2002). The key to the evolution of prey polymorphism in such system is that predators are better able to discriminate prey from the background when prey are frequently encountered. Thus, prey morphs found in low frequency in a population have a selective advantage. An analogous process emerged in our simulations, in which the receiver individual learned the appropriate response towards individuals with a common badge size in the population. When an individual has a rare badge size it can be favoured or unfavoured by the naive behaviour. When the naive behaviour is peaceful, polymorphism is promoted, whereas when the naive behaviour is aggressive polymorphism is prevented. These two situations show how behaviours dependent on learning processes seem to generally trigger frequency-dependent selection because the learning algorithm collects more information on the more frequent values of the the trait distribution. For simplicity, we modelled a single dimension (badge size), yet the frequency-dependent effect of learning could potentially be more important when individuals learn from multidimensional signals.

Originally, Rohwer (1975) proposed the idea that certain phenotypic traits could be used by animals as status signals or signals of quality. This idea has been tested repeatedly on the dark patches of some bird species, such as the bibs exhibited by species in various families, particularly sparrows. So, are bibs real signals of quality? If so, are they handicaps or badge of status? From an evolutionary perspective it is unclear how the stability of status signals can be maintained. One option is that honesty is maintained by the cost of signal production; that is, when the signal works as a handicap (Botero et al. 2010; Grafen 1990; Johnstone and Norris 1993). Alternatively, if the signal does not carry production costs inversely proportional to quality, then it may work as a convention. In this latter case, honesty is presumably maintained by the aggressive reaction of receivers when the convention is broken (Enquist, Ghirlanda, and Hurd 2010; Tibbetts and Dale 2004). Our model captures both kinds of signals, although with some nuances. When we assume the handicap principle, simulations result in the evolution of honest signals mediated by the learned response. When we assume a cost-free signal, under certain conditions the learned response mediated the emergence of phenotypic variation in the signal. This variation, particularly in the mid-range of the distribution, facilitates the establishment of a convention: in this range, individuals respond appropriately to the trait of their peer. A simple correlation between the size os a signal and dominance rank (and quality) is expected under the handicap principle, as it is shown here and elsewhere (Botero et al. 2010; Grafen 1990; Johnstone and Norris 1993). However, the branching event we documented shows that conventions can emerge in a less straightforward way when mediated by learning processes. These nuances could potentially make sense of seemingly contradictory evidence on the correlation between plumage traits and quality. A case in point is that of the House Sparrow (*Passer domesticus*), a textbook example of badges of status. According to the prevailing narrative (Senar 2006; Searcy and Nowicki 2005), the bib size of males of this species is a signal of dominance rank and quality. However, a recent meta-analysis called into question this narrative by showing that the effect size of the association between bib size and dominance rank is small and uncertain (Sánchez-Tójar et al. 2018). Our simulations revealed that, conventions arose only in an intermediate range of the signalling trait, yet empirical studies seeing for correlations between phenotypic traits and quality are often performed across the whole range of variation. Whether such a situation may apply empirically to bib sizes in House Sparrows remains to be studied.

The focus of theoretical and empirical work on communication systems, and particularly badges of status, is often explaining the presence and absence of certain morphological traits within populations. Phenotypic traits hypothesized to play a role in communication systems regularly exhibit patterns of geographic variation within species that have long called for explanations. One example of this is the leapfrog pattern (Remsen 1984), whereby a certain morphological trait alternates its presence and absence in a set of geographically adjunct populations, likely as a consequence of selection (Cadena, Cheviron, and Funk 2011; Márquez et al. 2020). However, it is unclear what type of ecological process is behind these selective regimes. Given the wide variation in outcomes found in our model, which depend on both stochasticity and cognitive parameters which may vary in space owing to various processes, we speculate that variation in cognition and learning could provide some explanatory power in this respect. A first step to assess this hypothesis would be to evaluate the way individuals in different populations respond to novel traits (*e.g.* Avendaño and Cadena 2021).

In sum, we have presented here an evolutionary model where the evolution of a communication system is mediated through the learned responses of receivers. This is a novel way to understand the evolution of communication that integrates classical cognitive process in learning with evolutionary explanations of communication. This approach contributes to the further integration of proximate and ultimate explanations in behavioral and evolutionary biology.

---

**Figure S1:**
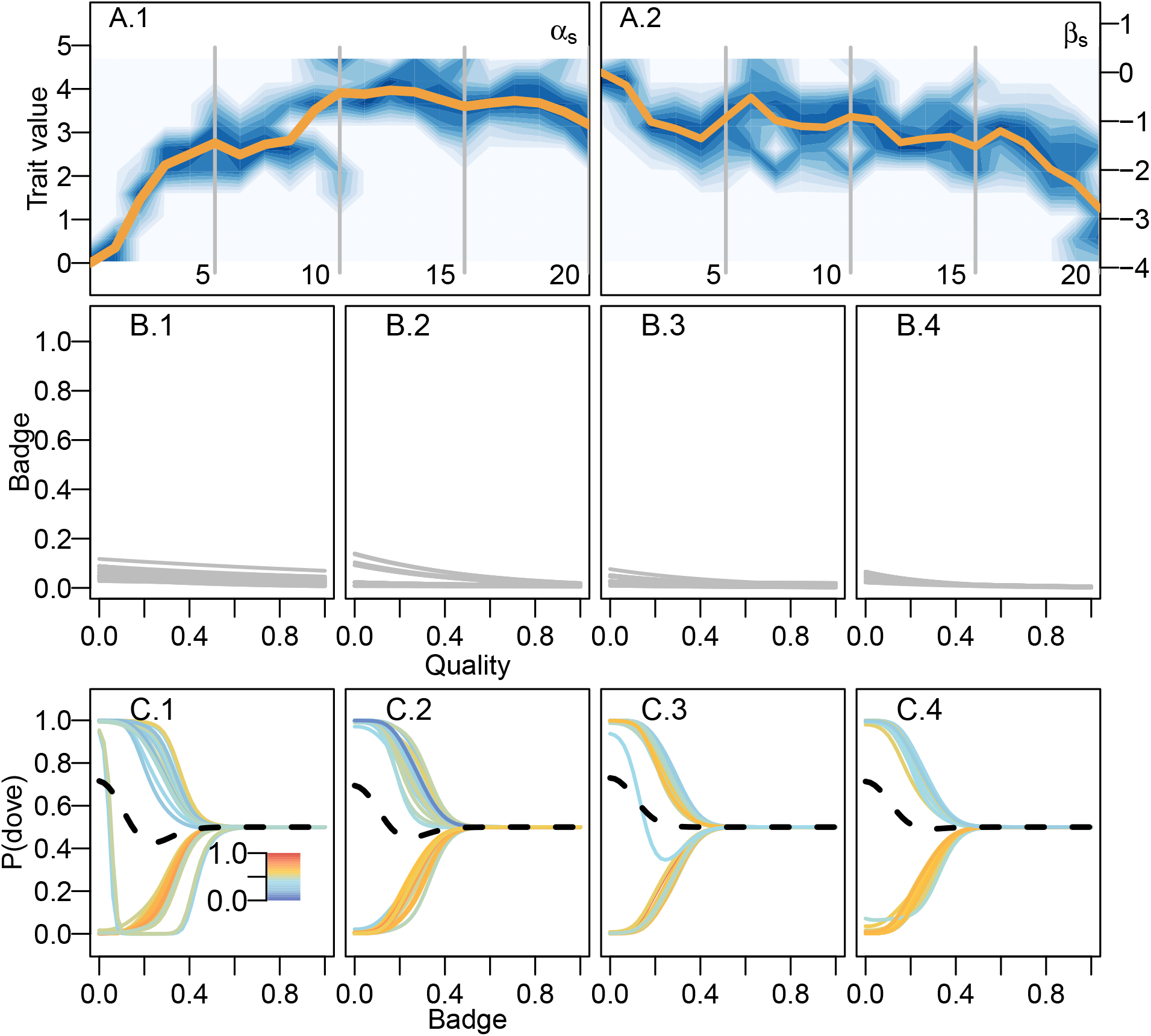
The evolution of cheap signals. Portrait of the evolutionary dynamics of the sender code with snapsots of both sender and receiver codes just as in fig. 2. The middle panels show changes in the distribution of values for α and β along evolutionary time. Panels above and below correspond to snapshots of the sender and receiver codes, respectively, generation time of the snapshots are indicated by the grey lines in the middle panels.

**Figure S2:**
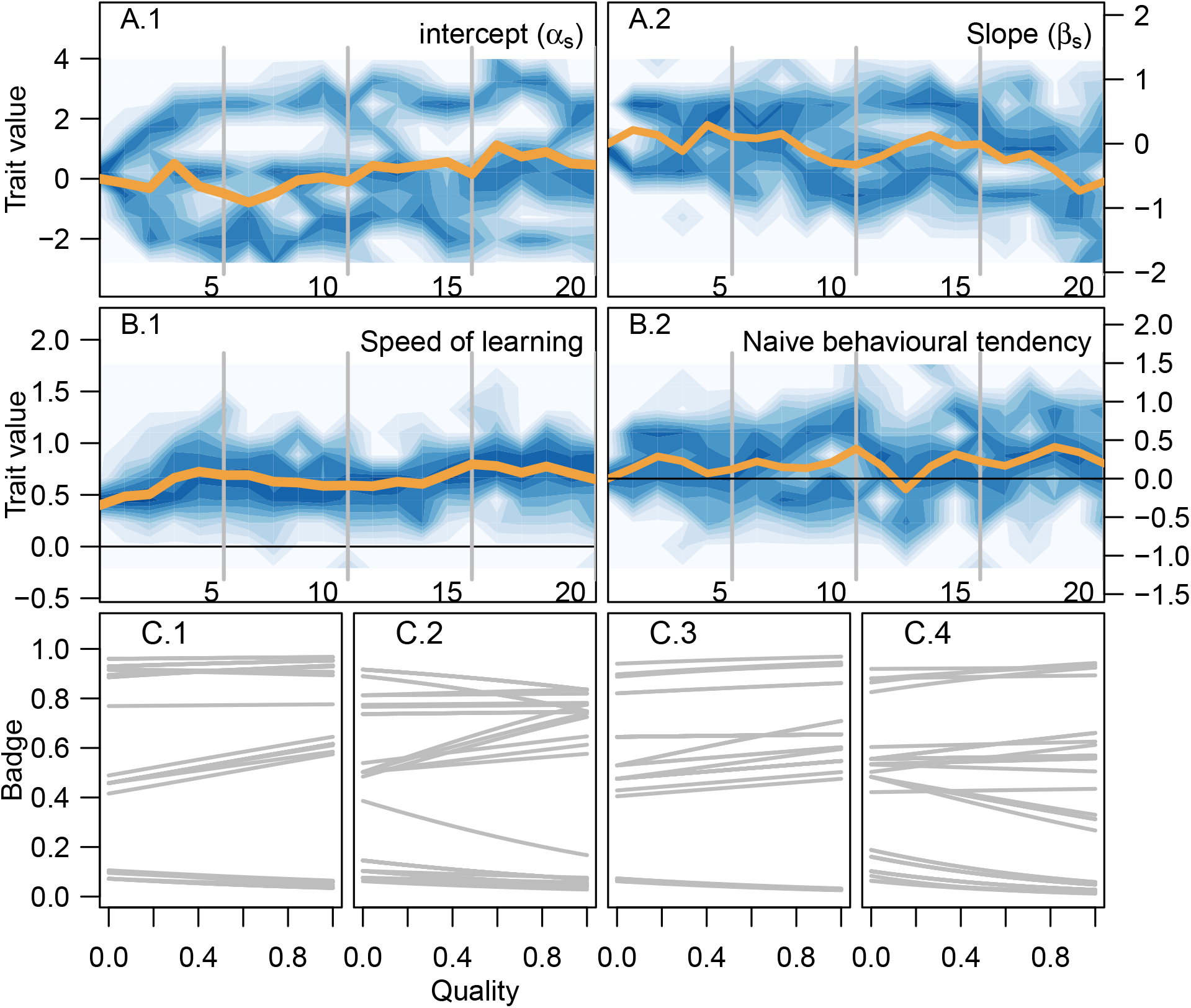
Coevolutionary dynamics of both the signal reaction norm and key parameters of the learning module. Dynamics are portrait as changes in the distribution of values assorting in the population. In the top panels (A) evolutionary dynamics of the sender code; A.1 shows the intercept and A.2 the slope of the logistic reaction norm. Both of these parameters experience evolutionary branching, whereby in the end three distincs types are assorting in the population. The middle panels (B) show changes in the distribution of values for the speed of learning (*A*;B1) and the behavioural tendency before learning (*y*^0^; B2) along evolutionary time. The speed of learning does not change drastically, it maintains values well above 0 (black line). In contrast, the initial behavioural tendency does not evolve away from 0 (black line). Both of these conditions favour evolutionary branching in the signal reaction norm. Bottom Panels correspond to snapshots of the sender code. Generation time of the snapshots are indicated by the grey lines in the top and middle panels.

**Table S1:** Notation of the model parameters and variables.

| Symbol | Description |
| --- | --- |
| $Q_i$ | Quality of individual $i$ |
| $N$ | Population size |
| $n$ | Number of interactions as the focal individual |
| $\sigma$ | Standard deviation of the truncated normal distribution from which quality is drawn |
| $B_i$ | Badge size of individual $i$ |
| $\alpha_i$ | Intercept of the badge size reaction norm for individual $i$ |
| $\beta_i$ | Slope of the badge size reaction norm for individual $i$ |
| $\mu$ | Mutation rate |
| $\sigma_\mu$ | Standard deviation of the normal distribution from which quality is mutations are drawn |
| $s_i$ | Survival probability of individual $i$ |
| $k_1$ | Intercept of the survival function |
| $k_2$ | Slope of the survival function |
| $V$ | Value of the contested resource |
| $C$ | Cost of an escalated fight |
| $p_{ij}$ | Probability that individual $i$ wins an escalated fight against individual $j$ |
| $k_3$ | Parameter determining how important is quality defining the probability of wining |
| $\delta$ | Prediction error |
| $A$ | Speed of learning |
| $c$ | Number of feature centres |
| $b_z$ | badge size for feature centre $z$ |
| $\hat{R}$ | Reward estimate |
| $x_z$ | weight of feature centre $z$ on the reward estimation |
| $\theta$ | width of the generalization function |
| $q_{ij}$ | probability that individual $i$ retreats from a fight in an encounter with individual $j$ |
| $y_z$ | weight of feature centre $z$ on the tendency to retreat |
| $w_i$ | total pay-off of individual $i$ |
| $w_0$ | base line pay-off |

